# NOVA2 regulates neural circRNA biogenesis

**DOI:** 10.1101/2021.05.02.442201

**Authors:** David Knupp, Daphne A. Cooper, Yuhki Saito, Robert B. Darnell, Pedro Miura

**Affiliations:** Department of Biology, University of Nevada, Reno, Reno, NV 89557, USA; Laboratory of Molecular Neuro-oncology and Howard Hughes Medical Institute, The Rockefeller University, New York, NY 10065, USA

## Abstract

Circular RNAs (circRNAs) are highly expressed in the brain and their expression increases during neuronal differentiation. The factors regulating circRNAs in the developing mouse brain are unknown. NOVA1 and NOVA2 are neural-enriched RNA-binding proteins with well-characterized roles in alternative splicing. Profiling of circRNAs from RNA-seq data revealed that global circRNA levels were reduced in embryonic cortex of *Nova2* but not *Nova1* knockout mice. Analysis of isolated inhibitory and excitatory cortical neurons lacking NOVA2 revealed an even more dramatic reduction of circRNAs and establish a widespread role for NOVA2 in enhancing circRNA biogenesis. To investigate the *cis*-elements controlling NOVA2-regulation of circRNA biogenesis, we generated a backsplicing reporter based on the *Efnb2* gene. We found that NOVA2-mediated backsplicing of circ-*Efnb2* was impaired when YCAY clusters located in flanking introns were removed. CLIP and additional reporter analysis demonstrated the importance of NOVA2 binding sites located in both flanking introns of circRNA loci. NOVA2 is the first RNA-binding protein identified to globally promote circRNA biogenesis in the developing brain.

## INTRODUCTION

Alternative splicing (AS) affects approximately 95% of human multi-exon genes (1). Through AS, hundreds of thousands of RNA isoforms with distinct structural properties, localization patterns, and translation efficiencies can be expressed as protein isoforms with diverse functions (2). In the mammalian nervous system, AS is especially pervasive and highly conserved (3,4). During brain development, AS is responsible for establishing neuron-specific splicing patterns at defined stages, and developmentally regulated alternative exons have essential roles in synapse formation, neuronal migration and axon guidance (5–8). Stage-specific AS patterns during development are controlled by RNA binding proteins (RBPs) enriched or specifically expressed in neurons and are critical for proper development as their dysregulation underlies many neurological disorders (9–13).

Circular RNAs (circRNAs) are generated through backsplicing, a type of AS (14). During backsplicing, the downstream 5’ splice site (SS) covalently bonds the upstream 3’ SS of a circularizing exon creating a closed loop “circle” that is resistant to exoribonuclease digestion (14). Most characterized circRNAs are derived from annotated exons of protein coding genes, and many are independently regulated from their host genes and exhibit unique expression patterns over various time-points (15). Thus far, ascribed circRNA functions include sequestration of microRNAs, translation of small peptides, modulation of the immune response, and transportation and scaffolding of RBPs (16–26). CircRNAs are enriched in brain tissues on a genome-wide level (27–30) and are dramatically upregulated during neural differentiation and maturation (15,29,30). CircRNAs are also found to accumulate during aging and this trend appears to be specific to brain tissues and neurons (31–33).The functional significance of brain-expressed circRNAs is emerging, with only a handful of circRNAs found to have roles in the nervous system (34–36).

Several, RBPs have been identified to regulate circRNA biogenesis, including Muscleblind, Quaking (QKI), ADAR, FUS, and several hnRNPs and SR proteins (37–41). However, investigation of circRNA regulation by RBPs from *in vivo* brain or neuron datasets is lacking. There are several well-characterized splicing factors with roles in the nervous system including RBFOX1/2/3, PTPBP1/2, nrSR100/Srrm4, Hu proteins, and NOVA1/2 (8). Deletion or dysregulation of any of these splicing factors results in serious and often lethal neurological defects (8). Among the best characterized neural-enriched splicing factors are the NOVA proteins, which were originally discovered as autoantigens in patients with paraneoplastic opsoclonus-myoclonus ataxia, a neurological condition characterized by motor and cognitive defects (42–44). NOVA1 and NOVA2 are paralogues that bind clusters of YCAY motifs in single-stranded RNA hairpin loops to regulate alternative splicing (45–47). Knockout (KO) of either paralogue in mice results in early lethality. This has been attributed to death of motor neurons in the case of NOVA1 deficiency, and aberrant migration of cortical and Purkinje neurons in the case of NOVA2 (13,48-50). Although both proteins recognize the same RNA binding motif, their expression is largely reciprocal. For instance, immunohistochemical and *in situ* hybridization data indicate NOVA2 is lowly expressed in midbrain and spinal cord while high expression is observed in the cortex and hippocampus. In contrast, NOVA1 is highly expressed in the midbrain and spinal cord while relatively lowly expressed in the cortex (13,44). Furthermore, NOVA1 and NOVA2 appear to have different splicing regulatory networks in the developing cortex itself (13). More recently, conditional knockout (cKO) of NOVA2 in either excitatory (Emx1+) or inhibitory (Gad2+) neurons led to thousands of AS events that were largely unique to each cell-type (50). In addition, loss of NOVA2 in excitatory neurons resulted in disorganization of cortical and hippocampal laminar structures, whereas this was not observed in NOVA2-KO inhibitory neurons. These findings showcase the importance of examining AS outcomes among different neuron subtypes.

In this study, we present a novel role for NOVA2 as a regulator of backsplicing in mouse cortical neurons. Global circRNA analysis of RNA-seq data revealed that loss of NOVA2 but not NOVA1 caused an overall decrease in circRNA expression. To investigate NOVA2 regulation of circRNAs specifically in neurons, we examined sorted cortical excitatory and inhibitory neuron subpopulations lacking NOVA2. We found that loss of NOVA2 in either subpopulation led to a dramatic reduction of circRNA expression and that this trend was independent of host gene expression. We identified one highly conserved and abundant circRNA, circ*Efnb2*, that was strongly downregulated in all NOVA2-deficient conditions. In addition, we investigate the role and positional importance of NOVA2 intronic binding sites for the regulation of circRNA backsplicing.

## MATERIALS AND METHODS

### Accession numbers

Wild-Type (WT), *Nova1-KO* and *Nova2-KO* whole cortex RNA-seq data were obtained from GEO under accession number GSE69711. FACs sorted *Nova2*-KO neuron RNA-seq datasets were obtained from GEO under accession number GSE103316. For individual accession numbers for each see **Supplementary File S1**.

### Mouse tissue preparation, RNA extraction, RNaseR Treatment, and RT-qPCR

All procedures in mice were performed in compliance with protocols approved by the Institutional Animal Care and Use Committee (IACUC) of the Rockefeller University and the University of Nevada, Reno. Mouse tissue samples were pulverized using a mortar and pestle on dry ice, and RNA was extracted using Trizol (ThermoFisher Scientific). For harvesting cultured cells PBS washes were performed followed by Trizol extraction. For RNase R treatment, 100μg total RNA from cortex or cultured HEK293 cells was treated with or without 1μL of RNaseR [20 U/μl] (Lucigen), plus 1.9μL RNaseOUT [40 U/μL] (ThermoFisher Scientific) and 1μL Turbo-DNase [2 U/μL] (Invitrogen) in a 60μL reaction volume for 30 minutes at 37°C. RNase R reactions were terminated and RNA was purified as previously described (51). Equal amounts of RNase R or mock treated RNA served as input for cDNA preparation. PolyA+ RNA and polyA- RNA was obtained using previously described methods (51). Briefly, the NucleoTrap mRNA column-based kit (Machery-Nagel) was used according to the manufacturers protocol. RNA present in the flow-through (not bound to oligo(dT) beads) was precipitated to isolate polyA-depleted RNA fraction. For RT-qPCR experiments, Turbo-DNase (Invitrogen) treated RNA was reverse transcribed using random hexamers (Invitrogen) and Maxima reverse transcriptase (ThermoFisher Scientific) according to the manufacturer’s specifications. RT-qPCR was performed on a BioRad CFX96 real time PCR machine using SYBR select mastermix for CFX (Applied Biosystems). The delta delta Ct method was used for quantification. Target gene expression for both circRNA and host-mRNA expression was normalized to *Gapdh*. Experiments were performed using biological triplicates. Student’s t-test (two-tailed and unpaired) was used to test for statistical significance.

### Cell Culture

HEK293 cells (ATCC) were cultured in DMEM (ThermoFisher Scientific) with 10% FBS (Atlanta Biologicals). HEK293 cells were transfected with NOVA2 expression plasmid or empty vector control (Gifts from Dr. Zhe Chen at University of Minnesota) in addition to the circ*Efnb2* or circ*Min*i backsplicing reporters, using PEI transfection reagent (Polysciences), and reduced serum medium Opti-MEM (ThermoFisher Scientific). The cells were cultured for 24 hours prior to RNA extraction.

### Northern Blotting

RNA samples were denatured by mixing with 3 volumes NorthernMax Formaldehyde loading dye (ThermoFisher Scientific) and ethidium bromide at a final concentration of 10 μg/mL for 15min at 65°C. Denatured samples were loaded onto a 1% MOPS gel with 1x Denaturing Gel Buffer (ThermoFisher Scientific) and ran at 102V for 60 min. RNA samples were transferred to a Cytiva Whatman Nytran SuperCharge membrane (ThermoFisher Scientific) for 1.5 hours using a Whatman TurboBlotter transfer system (ThermoFisher Scientific) and NorthernMax Transfer Buffer. Samples were then UV cross-linked with a Stratagene linker to the nylon membrane prior to probe hybridization. Double-stranded DNA probes were prepared by end-point PCR and labeled with dCTP [α-32P] (PerkinElmer) using the Cytvia Amersham Megaprimer labeling kit (Thermo Fisher Scientific) according to manufacturer’s instructions. Blots were hybridized overnight in ULTRAhyb™ Ultrasensitive Hybridization buffer (ThermoFisher Scientific) at 42°C. Following hybridization, blots were washed at room temperature 2×5 min in a low-stringency buffer followed by 2×15 min in a high-stringency buffer at 42°C. Blots were then exposed for 4-5 days to a GE Storage Phosphor screen (Millipore Sigma) before imaging on a Typhoon™ FLA 7000 imager (GE). Probe sequences are listed in **Supplementary File S9**.

### RNA-Seq Analysis for CircRNA prediction, mapping and differential expression

For de novo identification of circRNAs a custom pipeline was carried out. First, raw FASTQ files were aligned to the mm9 genome using HISAT2 v2.1.0 (52) (parameters: --no-mixed –no-discordant). Unmapped reads were aligned using BWA v0.7.8-r455 mem and CIRI2 v2.0.4 (default parameters) was used to obtain a set of predicted circRNA loci. For alignment to circRNA junction spanning FASTA sequence templates of 220 nt we used Bowtie2 v2.2.5 (53) (parameters: -score-min=C,−15,0). After mapping, Picard (http://broadinstitute.github.io/picard) (parameters: MarkDuplicates ASSUME_SORTED=true REMOVE_DUPLICATES=true) was used to remove duplicates from our mapping data. To quantify the number of mapped reads for each junction we used featurecounts v1.5.0 (54) (parameters: -C -t circle_junction).

For each dataset (*Nova1*-KO vs. WT, *Nova2*-KO vs. WT, *Nova2*-cKO^tdTomato;Emx1-Cre^ vs. WT, *Nova2*-cKO^tdTomato;Gad2-Cre^ vs. WT) a cutoff of 6 reads across the 6 libraries for each condition (3 biological replicates per condition) was set. To account for difference in library depth among samples, scaling by circRNA Counts Per Million of reads (CPM) in each library was performed. Fold-change CPM values were generated between KO and WT conditions. A cutoff of 2.0-fold-change difference and significance threshold of *P*<0.05 via t-test was used across the normalized CPM values to identify differentially expressed circRNAs. Correction for multiple hypothesis testing was not performed.

The CircTest pipeline was performed to quantify host gene independent circRNA expression patterns. DCC/CircTest v0.4.7 was used to quantify host gene read counts. In accordance with DCC pipeline (55), FASTQ files were mapped with STAR 2.6.0b using the recommended parameters and aligned to the GRCm38 genome. The circRNA and linear RNA counts obtained from DCC were used as input for the Circ.test module (parameters: Nreplcates=3 filter.sample=4 filter.count =3 percentage = 0.1 circle_description=c(1:3)). To define a circRNA as host gene independently expressed, we used the default parameter, adj. *P*<0.05 (Benjamini-Hochberg correction). ggplot2 (56) R packages and custom scripts were used to generate all plots.

### Mapping and quantification of linear RNA expression

Reads were aligned to the NCBI37 reference genome using HISAT2 v2.1.0 (GENCODE annotations M1 release, NCBIM37, Ensembl 65). FeatureCounts v1.5.0 (parameters -t exon -g gene_id) was used to obtain a counts table as input for differential expression analysis. For differential expression analysis, DESeq2 v1.26.0 was performed using Benjamini-Hochberg correction and apeglm Bayesian shrinkage estimators with a 2.0-fold-change and adj. *P*<0.05 required to consider a linear RNA as differentially expressed. For alternative splicing analysis, the rMATS pipeline was used to calculate significant exon skipping events in *Nova2*-cKO^tdTomato;Emx1-Cre^ versus WT and *Nova2*-cKO^tdTomato;Gad2-Cre^ versus WT datasets. FASTQ files were mapped using STAR 2.6.0b using default parameters (parameters: -- chimSegmentMin 2 --outFilterMismatchNmax 3 –alignEndsType EndToEnd –runThreadN 4 – outSAMstrandField intronMotif –outSAMtype BAM SortedByCoordinate –alignSJDBoverhang 6 – alignIntronMax 30000) and aligned to the GRCm38/mm10 genome (GENCODE annotations vM22 release). rMATs-v.3.2.5 was used to discover significant alternative splice events under the default parameters. For post processing, we filtered the output file, SE.MATS.ReadsOnTargetAndJunctionCounts.txt to contain skipping events with FDR<0.01. Mapped circRNA BED files were converted to mm10 coordinates using UCSC genome browser liftover tool. Then bedtools suite was used to find overlap with significant exon skipping events. Overlaps were manually checked using Integrative Genomics Viewer (IGV) (57).

### Backsplicing Reporter Assays

All plasmids are available upon request. The pUC19 plasmid backbone was used to generate circ*Efnb2* and circ*Mini* backsplicing reporters with modifications. Briefly, the CMV enhancer/promoter region was amplified from the pcDNA3.0 backbone using Phusion High-Fidelity Polymerase (NEB) and subcloned into the pUC19 vector upstream of the circ*Efnb2* or circ*Mini* backsplicing cassettes. In addition, BGH and rB-Globin poly(A) sequences were subcloned downstream of the backsplicing cassettes using synthetic gene fragments (gBlocks, IDT) for transcription termination. All fragments used to generate the backbone vector and subsequent backsplicing cassettes were cloned using the NEB Hifi assembly kit (NEB) following the manufacturers protocols.

For circ*Efnb2* reporter three genomic fragments were amplified by PCR using Phusion High-Fidelity Polymerase. The three genomic fragments (mm10 coordinates) are as follows: 1) Truncated upstream *Efnb2* exon1 (81 bp) plus downstream flanking intron (134 bp) (chr8:8660350-8660564); 2) circularizing *Efnb2* exon 2 (284 bp) plus partial upstream (448 bp) and downstream (475 bp) flanking introns (chr8:8638731-8639937) and 3) truncated *Efnb2* exon 3 (72 bp) plus partial intronic upstream sequence (197 bp) (chr8:8623169-8623437); (chr8:8660350-8660564). Initial transfection experiments with circ*Efnb2*-WT demonstrated two unintended backspliced products originating from the AmpR cassette and non-coding sequence immediately downstream of the CMV promoter when examined by RT-PCR. As a result, we introduced two silent mutations into the AmpR coding sequence and deleted 115 bp of non-essential sequence between the CMV promoter and circ*Efnb2* cassette. Follow-up experiments showed all unintended backspliced products were abolished. Mutations located in the circ*Efnb2* introns were introduced by PCR amplification of the pWT backbone using Phusion High-Fidelity polymerase and ligation with gBlock donor DNA carrying point mutations targeting YCAY motifs. Mutations were confirmed by Sanger sequencing.

To generate the artificial circMini-WT vector, two gBlocks consisting of full-length GFP coding sequence, partial human *ZKscan1* intron sequence and partial *Mboat2* intron sequence were cloned into the modified pUC19 vector described above. We used the Berkeley Drosophila Genome Project splice prediction tool (https://www.fruitfly.org/seq_tools/splice.html) with default settings to guide GFP and intron sequence modifications that would improve splicing efficiency and prevent unintended splice products from being generated. In addition, existing YCAY motifs were mutated to prevent NOVA2 binding. Artificial 10x YCAY sequence was produced via gBlocks and were cloned 50 bp upstream (pMini-UP), 49 bp downstream (pMini-Down) or in both locations (pMini-Both) in relation to the circRNA loci.

## RESULTS

### Global circRNA levels are reduced in *Nova2*-KO whole cortex

To investigate potential factors that might contribute to regulation of circRNAs in the murine brain, we analyzed paired-end total RNA-seq data from embryonic *Nova1-KO* and *Nova2*-KO mouse cortex samples for changes in circRNA expression (13) (**Supplementary File S1**). These RNA-seq libraries were generated using random hexamer-based priming as opposed to oligo(dT) priming for cDNA synthesis, thus enabling detection of circRNAs which are not-polyadenylated. CircRNAs were identified by back-splice junction (BSJ) reads using the CIRI2 algorithm (**Figure 1A**). We set a minimum expression threshold of 6 BSJ reads across the 6 libraries (minimum average of 1 read per biological replicate) for each dataset, resulting in 1565 and 3708 exonic circRNAs identified for the *Nova1* and *Nova2* datasets, respectively (**Supplementary File S2 and File S3**). BSJ read counts were normalized to library size to obtain Counts Per Million mapped reads (CPM). Global circRNA CPM values were found to be significantly reduced in *Nova2*-KO samples compared to controls (*P*<2.48×10^−11^, Wilcoxon-rank sum test with continuity correction) (**Figure 1B**). In contrast, global circRNA levels were not altered in *Nova1*-KO mice compared to controls (**Figure 1B**).

**Figure 1.**
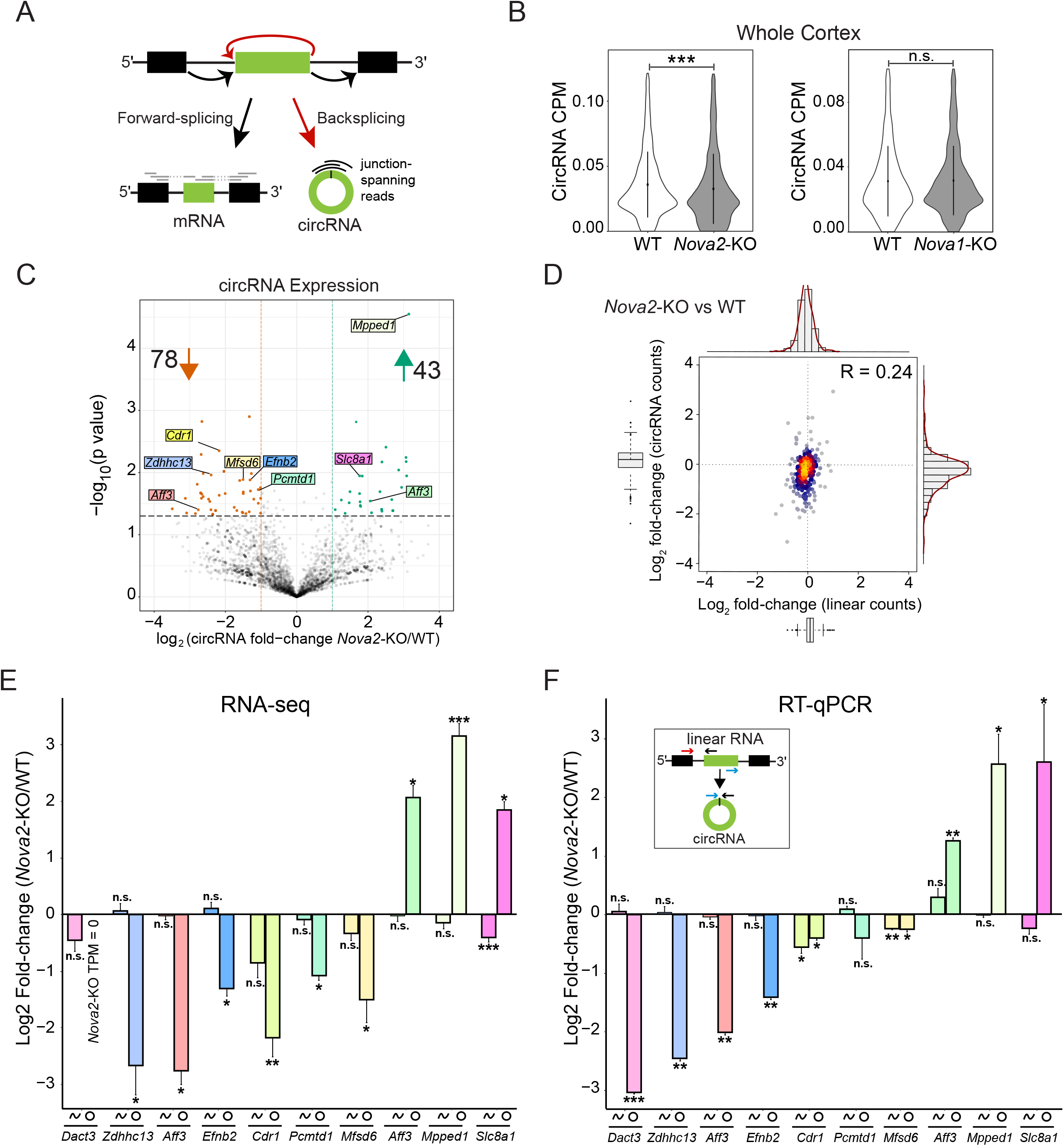
NOVA2 regulation of circRNA biogenesis in mouse cortex. (**A**) Schematic of forward-spliced and backspliced read alignments for detection of linear RNA and circRNA expression, respectively. The circularizing exon is shown in green. (**B**) CircRNA CPM is significantly reduced in *Nova2-*KO (*left*) but not *Nova1-*KO (*right*) whole cortex. Significance reflects non-parametrical Wilcoxon rank-sum test with continuity correction. n = 3 biological replicates for each condition. (**C**) Volcano plot of circRNAs in *Nova2-*KO vs. WT showing circRNAs downregulated (orange dots) and upregulated (green dots) in the knockout condition (log_2_FC>1, *P*<0.05). (**D**) High density scatterplot of 311 circRNAs (minimum 3 BSJ read counts in 4 out of 6 replicates). Y-axis reflects log_2_ fold-change of circRNA counts. X-axis reflects log_2_ fold-change of linear counts from host genes. (**E**) RNA-seq expression of 10 circRNAs (7 downregulated, 3 upregulated) from *Nova2*-KO vs. WT and their corresponding host gene mRNA expression. (**F**) RT-qPCR validation of the same 10 regulated circRNAs and their host gene mRNAs normalized to *Gapdh*. Inset diagram depicts primer locations used for circRNA and host gene linear mRNA quantification. \**P*<0.05; \*\**P*<0.01; \*\*\**P*<0.001; n.s., not significant. Student’s t-test was used for statistical significance (two-tailed, unpaired). CPM, Counts Per Million.

In order to capture expression of individual circRNAs in *Nova2-*KO cortex we generated volcano plots using *P*-value and fold-change, as previously reported (32). We observed a slight trend for downregulation in *Nova2*-KO mouse cortex compared to WT samples. From the volcano plot, it is evident that more circRNAs were downregulated than upregulated in the KO condition (*P*<0.05, Log_2_FC>1) (**Figure 1C**). In contrast, the *Nova1*-KO dataset lacked any biased expression trend (**Figure S1A**). The reduced circRNA levels in *Nova2*-KO samples could have been a consequence of reduced transcriptional activity from the host genes that the circRNAs are derived from. Thus, we performed differential expression analysis for the mRNAs generated from the host genes of the regulated circRNAs (**Figure S1B**). Alignment was performed using HISAT2 (58), and DESeq2 (52) was used to perform differential expression analysis of mRNAs. No significant changes in host gene mRNA expression were detected. In addition, density plots were generated to contrast total read counts from circRNAs versus their linear RNA counterpart from the same host gene, read counts were obtained using DCC (see Materials and Methods) (55). We observed a clear downward shift along the y-axis, reflecting reduced circRNA expression, while linear RNA expression along the x-axis centered near zero indicating only minor expression changes (**Figure 1D**). Together, these data suggest that NOVA2-mediated regulation of circRNA biogenesis is largely independent of host gene transcriptional expression changes.

In order to provide experimental support for the circRNA expression trends, we performed RT-qPCR confirmation for 10 circRNAs that were either reduced (7 loci) or increased (3 loci) in the *Nova2*-KO condition (**Figure 1F**). For circRNA quantification, outward facing primers that only detect the circularized exons were used (**Figure 1F**). For quantification of the cognate mRNA, we employed primer sets with one primer located in an exon that is circularized and the other is located in the flanking upstream or downstream exon (**Figure 1F and Supplementary File S9**). Overall, we observed a good correlation between RNA-seq expression trends and our RT-qPCR results, confirming expression trends for 9/10 circRNAs and 8/10 host gene linear RNAs. In addition, we validated the circularity of these targets with RNase R, a 3’ to 5’ exoribonuclease that degrades linear RNAs while circRNAs are relatively more resistant (59). We found all 10 to be resistant to RNase R treatment, whereas the linear control gene, *Psmd4*, was degraded (**Figure S1C**). These experiments indicate that our sequencing analysis pipeline can detect bonafide circRNA expression changes.

### Loss of NOVA2 dramatically reduces global circRNA levels in isolated neuron subpopulations

NOVA2 expression is mostly limited to neurons (44). In contrast, circRNAs are expressed in various brain cell-types such as astrocytes, neurons, glia and oligodendrocytes (29,60). We reasoned that our analysis in whole cortex might obscure the specific regulation of circRNAs in neurons by NOVA2. Thus, we analyzed circRNAs in NOVA2 deficient neuron subpopulation datasets. Total RNA-seq data from fluorescence-activated cell sorted (FACS) embryonic inhibitory and excitatory cortical neurons deficient in NOVA2 (50) were analyzed using the CIRI2 pipeline. We identified 4123 and 2440 exonic circRNAs in Gad2+ and Emx1+ datasets, respectively, that passed our minimum BSJ read threshold (**Supplementary File S4 and S5**). Global circRNA levels were significantly decreased in both *Nova2*-KO datasets (*P*<2.2×10^−16^, Wilcoxon-rank sum test with continuity correction) (**Figure 2A, B**; inset violin plots). Similar to results from whole cortex, linear expression from the host gene of the differentially expressed circRNAs did not show any significant changes (**Figure S2A, B**). Volcano plots demonstrated a striking downregulation trend for hundreds of circRNAs in both inhibitory and excitatory neurons in the absence of NOVA2, with only a handful of upregulated circRNAs (**Figure 2A, B**). At least 9-fold more circRNAs were downregulated in either dataset compared to the number of circRNAs upregulated. Reduced circRNA expression in the knockouts remained when the analysis was performed with increased minimum thresholds of 10, 20, and 30 backspliced reads per condition (**Figure S3A-C**). Our results indicate that NOVA2 generally promotes circRNA biogenesis in cortical neurons.

**Figure 2.**
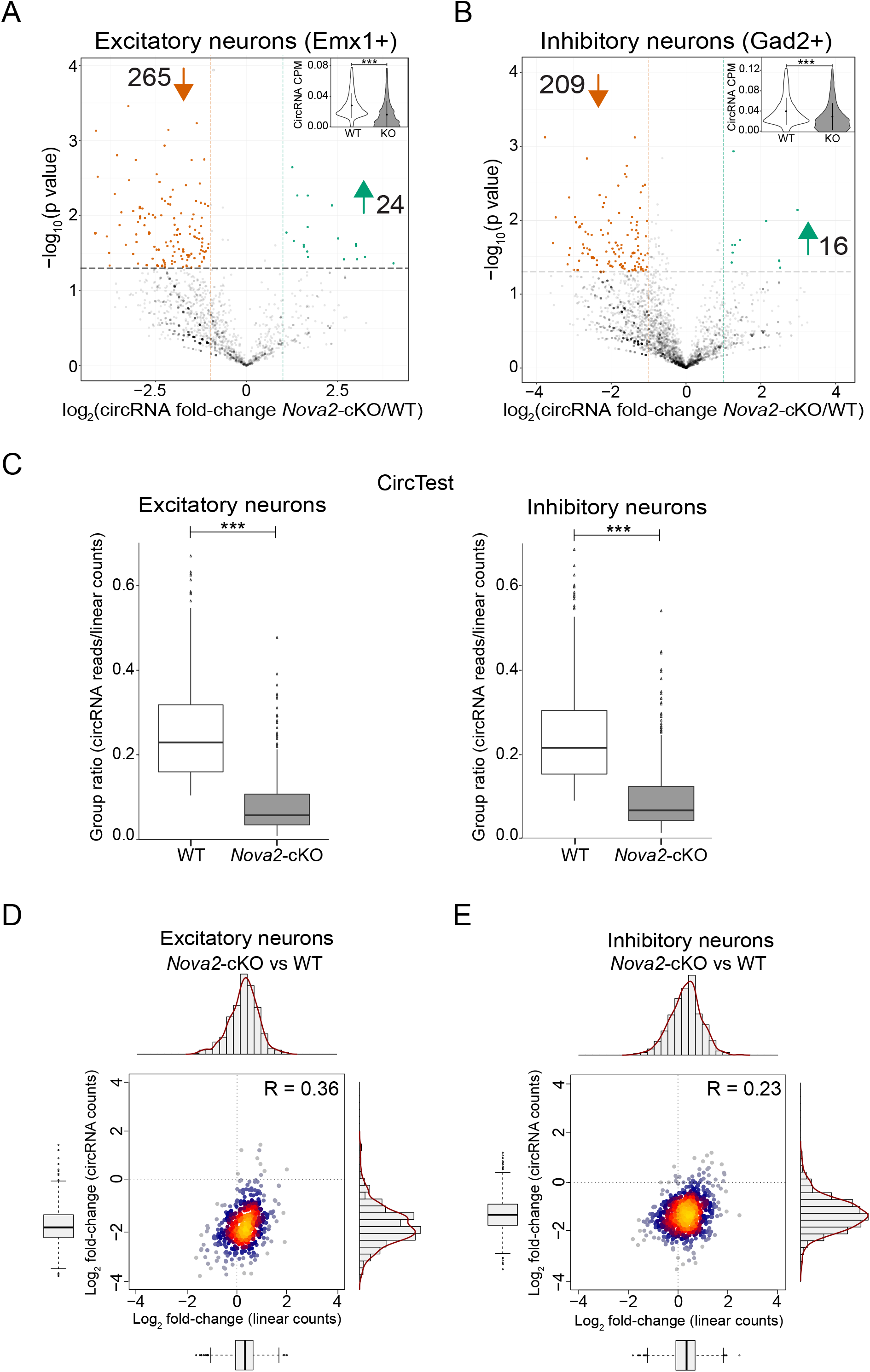
NOVA2 regulation of circRNA biogenesis in cortical excitatory and inhibitory neurons. (**A**) Volcano plot of circRNAs detected using CIRI2 in excitatory cortical neurons (Emx1+) and (**B**) inhibitory cortical neurons (Gad2+) deficient in NOVA2. In both datasets, more circRNAs were significantly downregulated compared to upregulated in *Nova2*-cKO cells relative to WT (log_2_FC>1, *P*<0.05). Inset violin plots show significant reduction in total circRNA CPM. Statistical analyses were carried out as in Figure 1B. (**C**) CircTest group ratio is significantly reduced in *Nova2-c*KO condition. Significance reflects non-parametrical Wilcoxon rank-sum test with continuity correction. n = 3 biological replicates for each condition. Y-axis; group ratio defined as the number of BSJ reads divided by the number linear-spliced reads. (**D**) High density scatterplot of 456 DCC/CircTest identified high confidence circRNAs (minimum 3 BSJ read counts in 4 out of 6 biological replicates) from excitatory neurons deficient in NOVA2. Y-axis reflects log_2_ fold-change of circRNA counts. X-axis reflects log_2_ fold-change of linear counts from host genes. Pearson correlation coefficient is shown in the upper right corner, indicating weak correlation between circRNA and linear RNA counts in *Nova2*-null excitatory neurons. (**E**) Same analysis repeated for *Nova2*-cKO inhibitory neurons with 495 high confidence circRNAs represented. \*\*\**P*<0.001. CPM, Counts Per Million.

We next analyzed the same datasets using DCC/CircTest, an independent circRNA analysis pipeline (55). DCC computes forward and backspliced read counts, while the CircTest module calculates the ratio of BSJ reads to linear, forward spliced reads and robustly tests for their independence. Applying a stringent filtering method (minimum 3 BSJ counts in 4 out of 6 biological replicates), we identified 519 and 750 circRNAs from excitatory and inhibitory cortical neuron populations, respectively (**Supplementary File S6**). In agreement with our CIRI2 analysis (**Figure 2A, B**), in the absence of NOVA2, we found a significant reduction of circRNA expression when normalized to the linear reads arising from the same host-gene (*P*<2.2×10^−16^, Wilcoxon-rank sum test with continuity correction; **Figure 2C**). Density plots showed a pronounced reduction in circRNA expression in both *Nova2*-KO neuron populations (vertical axis, **Figure 2D, E**). In contrast, linear RNAs from the same host gene showed only a minor shift to the right along the x-axis. Overall, we observed a weak correlation (Emx1+; R=0.36 and Gad2+; R=0.23) between circular and linear expression changes. We found that 456/519 Emx1+ expressed circRNAs (87%) and 495/750 of Gad2+ circRNAs (66%) passed CircTest significance testing for independence of circRNA and linear RNA expression (**Supplementary File S6**). Taken together, these results demonstrate that NOVA2-regulation of backsplicing is independent of linear host gene expression.

### NOVA2-regulated circRNAs and exon skipping events show little overlap

There are some reported instances of circRNA loci overlapping with exon skipping events (reviewed in (61)). We thus determined to what degree NOVA2 regulated exon skipping events (SE) overlapped with NOVA2 regulated circRNAs. We applied replicate Multivariate Analysis of Transcript Splicing (rMATS) (62) to probe for statistically significant SE events in the excitatory and inhibitory neuron datasets (**Supplementary File S7**). Returning to our CIRI2 generated list of differentially expressed circRNAs, we found that in excitatory neurons, only 3/24 (13%) upregulated circRNAs and 10/265 (4%) downregulated circRNAs overlapped with at least one significant SE event. Likewise, in inhibitory neurons only 1/16 (6%) upregulated circRNAs and 7/209 (3%) downregulated circRNAs overlapped with at least one significant SE event. Of note, all the circRNAs that overlapped with at least one SE event were multi-exonic, and in all instances only some of the exons within the circRNA loci were skipped. i.e., none of the regulated skipping events skipped an entire NOVA2-regulated circRNA. Given these results, we conclude that NOVA2-regulation of circRNAs is unrelated to NOVA2-mediated exon skipping.

### NOVA2-regulated circRNAs display cell-type specific regulation

We examined the overlap of NOVA2-regulated circRNAs between excitatory and inhibitory cell populations. We found that 247/293 and 120/225 of the NOVA2-regulated circRNAs in excitatory and inhibitory neurons, respectively, were expressed in both neuronal subtypes. Despite this broad overlap, we found that the identity of NOVA2-regulated circRNAs were largely distinct between the two cell-types (**Figure S4A**). Only 18 circRNAs were found to be NOVA2-regulated in both excitatory and inhibitory neurons. This is in line with what was previously found with the same datasets for linear alternative splicing (50). Thus, it appears that NOVA2 circRNA regulation exhibits neuronal sub-type specificity.

### Circ*Efnb2* is an abundant, conserved circRNA regulated by NOVA2

Having uncovered a genome-wide role for NOVA2 in circRNA regulation, we next turned to a single circRNAs locus for investigation into the mechanism. To choose a candidate for further study, we examined the differentially expressed circRNAs identified in both pipelines (CIRI2 or CircTest) and found that 74 in the Emx1^+^ dataset and 36 in the Gad2^+^ dataset overlapped (**Supplementary File S8**). Of these, only 7 circRNAs (all with reduced expression in *Nova2*-KO condition) were common to both Emx1 and Gad2 datasets (**Supplementary File S8**). This list included circ*0015034* (referred to from here on as circ*Efnb2*). Circ*Efnb2* is a 284 nt long circRNA generated from the 2^nd^ exon of the *Efnb2* gene. *Efnb2* encodes ephrin-B2, a transmembrane ligand which mediates cell-to-cell communication via contact with adjacent Eph receptor (63). The same locus also produces a circRNA in humans (circBaseID; hsa_circ_0029247) with identical primary sequence. Finally, circ*Efnb2* is abundant, ranking in the top 15% of high confidence circRNAs with respect to circRNA to mRNA ratio (**Figure 3B**).

**Figure 3.**
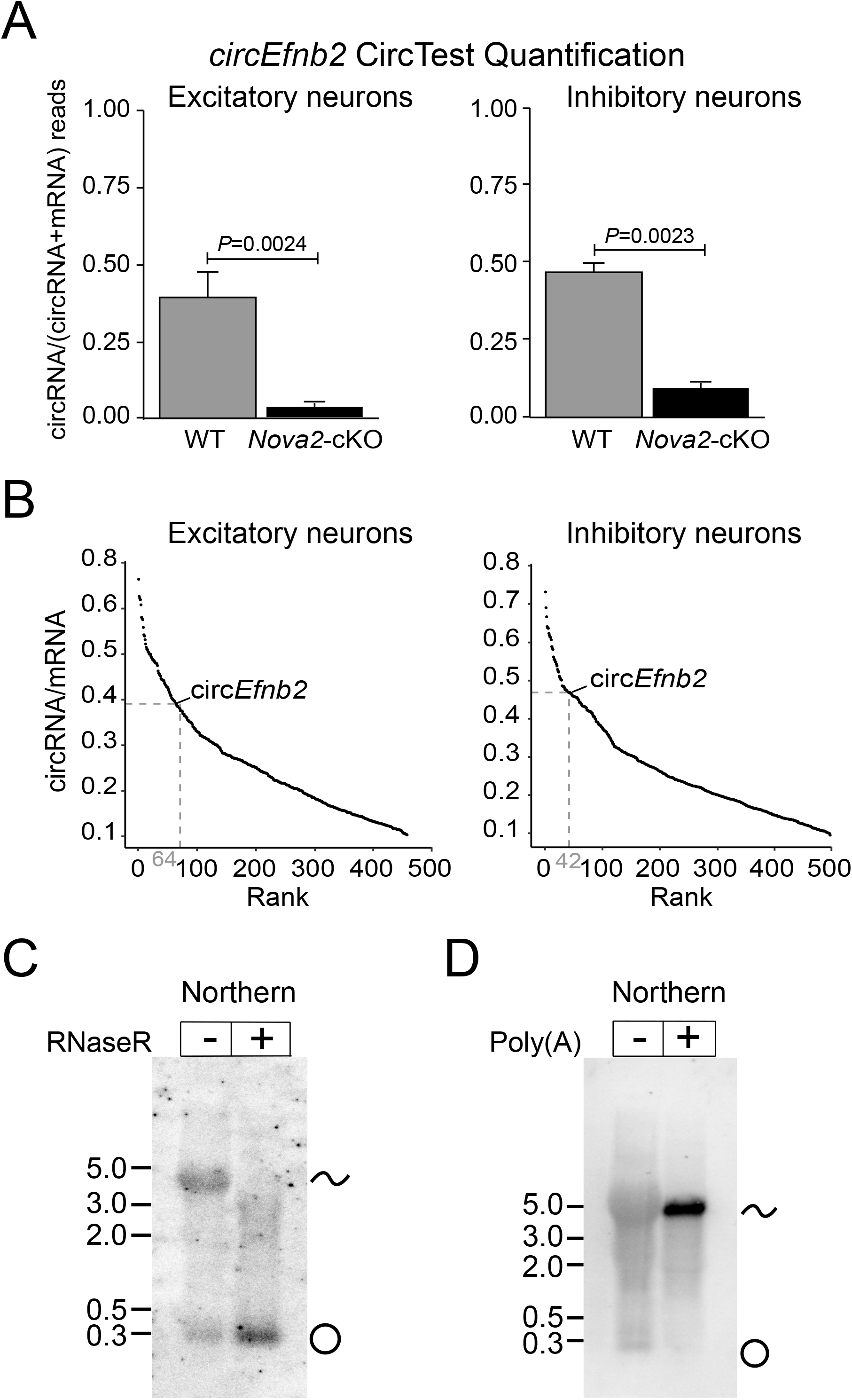
Circ*Efnb2* is an abundant, highly-conserved circRNA regulated by NOVA2. (**A**) CircRNA to mRNA ratio plot generated by CircTest. Ratio of circular junction read counts from circ*Efnb2* to average total counts at exon borders are shown. *P*-values were generated from CircTest module. (**B**) CircRNA/mRNA expression rank of *circEfnb2* in embryonic excitatory (64^th^) or inhibitory (42^nd^) neuron datasets (top 15%; both datasets). (**C**) Northern blot using probe overlapping circularized exon of *Efnb2* detects bands corresponding to the circRNA and mRNA from embryonic whole cortex RNA with or without RNase R treatment. (**D**) Northern blot performed for poly(A)- and poly(A)+ samples. RNA samples for Northern were obtained from E18 whole cortex.

To confirm the circularity of circ*Efnb2*, we performed Northern analysis of mouse cortex samples. We observed clear bands of the expected sizes for both the linear and circular products. Treatment with RNase R enriched circ*Efnb2* and depleted the linear transcript, confirming the circular and linear nature of the two major bands (**Figure 3C**). In addition, we captured polyA+ RNA from mouse cortex samples using oligo(dT) beads, as well as unbound RNAs (polyA- fraction) for Northern blot analysis. As expected, the polyA+ fraction enriched for polyadenylated, linear *Efnb2* mRNA and depleted the polyA tail-lacking circRNA (**Figure 3D**).

### Generation of a circEfnb2 backsplicing reporter

NOVA2 has a well-characterized YCAY binding motif (64). In order to determine which binding sites help facilitate NOVA2 regulation of *circEfnb2* we constructed a backsplicing reporter. To guide the design of the backsplicing reporters, we analyzed CLIP-peaks identified by publicly available NOVA2 CLIP-Seq datasets (50) (**Figure 4A**). Due to the extended lengths of the 5’ and 3’ flanking introns (20.9kb and 15.5kb, respectively), the circularizing exon combined with its full-length flanking introns is not amenable to plasmid subcloning and transfection. We thus opted to subclone truncated upstream and downstream flanking intronic regions that included major NOVA2-CLIP peaks (**Figure 4A**; “fragment 2”). In addition, we included partial sequences from exons 1 and 3 and ~150 bp of intron sequence to retain linear splicing from the reporter (**Figure 4A**; “fragment 1 and 3”). Fragment 3 also included prominent NOVA2-CLIP peaks. In the intronic sequences, we noted the presence of multiple YCAY motifs that were not associated with CLIP tags.

**Figure 4.**
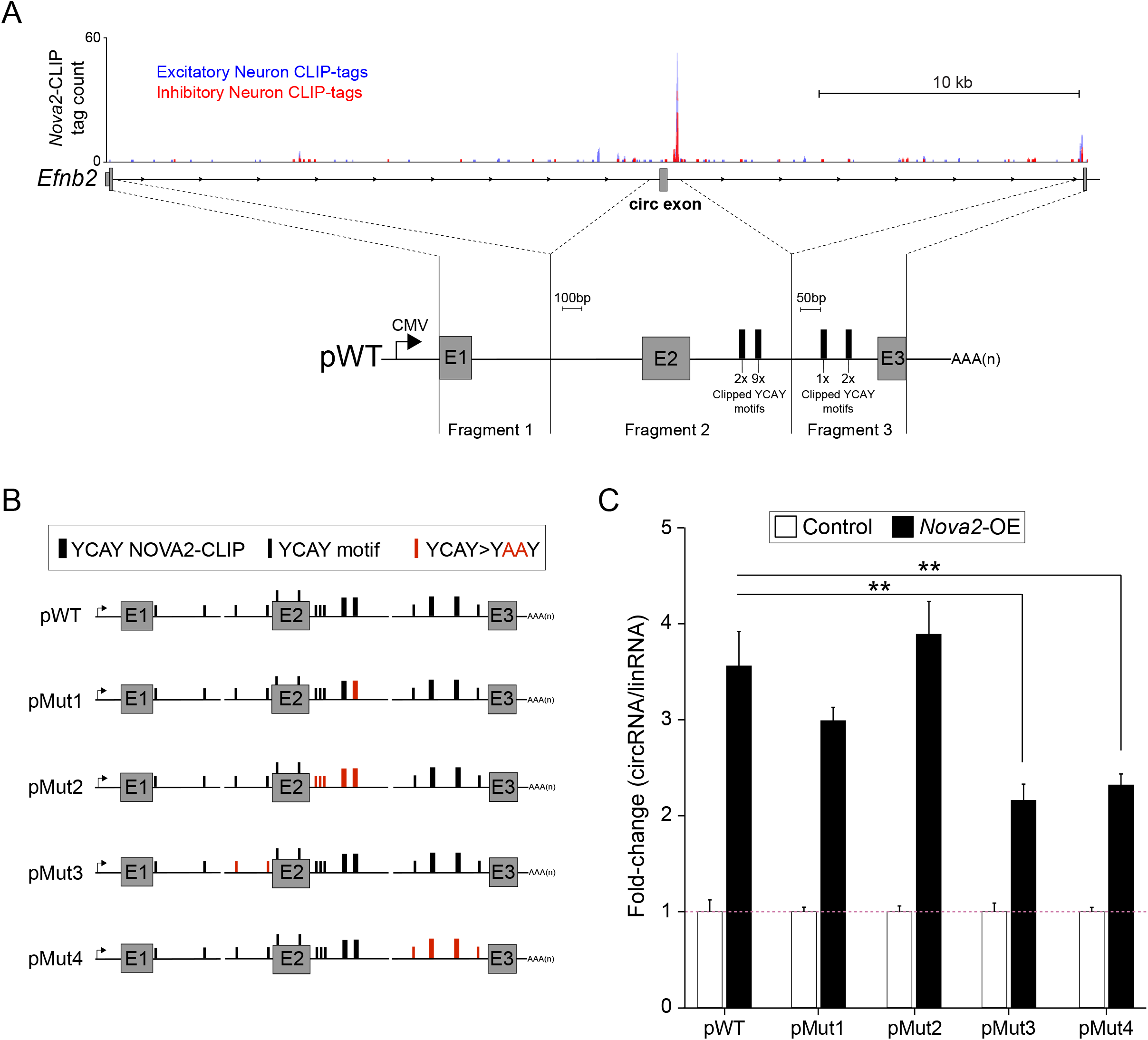
Role of intronic NOVA2 binding sites in Circ-*Efnb2* backsplicing. (**A**) Schematic of *Efnb2* locus shown in antisense (mm10, chr8:8623077-8660773). NOVA2-CLIP tags from excitatory cortical neurons (blue) or inhibitory neurons (red) that overlapped with NOVA2-CLIP peaks from either dataset were visualized using UCSC genome browser. Three genomic fragments used to construct the backsplicing reporter (pWT) are shown below, and the number of individual YCAY motifs present within each NOVA2-CLIP peak (thick black bar) are reported. Note, *Efnb2* Exon 1 (E1) and Exon 3 (E3) in the pWT backsplicing reporter are truncated, whereas circularizing Exon 2 (E2) is full length. (**B**) Reporter schematics for circEfnb2. NOVA2-CLIP peaks are represented by thick black bars as in panel A. YCAY motifs not associated with NOVA2-CLIP peaks are represented as thin black bars. Mutated YCAY motifs or peaks are shown in red. (**C**) RT-qPCR of circ*Efnb2* expression from reporters co-transfected with NOVA2 expression plasmid (*Nova2*-OE) or empty expression vector in HEK293 cells. For expression data, target genes were normalized to linear-spliced transcript (lin*Efnb2)* generated by the reporter. n=3 biological replicates. Error bars are represented as mean ± SEM. \*\**P*<0.01, compared to pWT circ*Efnb2* expression. (Students t-test, two-tailed and unpaired).

Consistent with previous NOVA2 splicing reporter studies, we examined regulation of circ*Efnb2* in HEK293 cells which express very low levels of *Nova2* (**Figure S5A**) (13,65–68). Initial analysis of the RNAs generated from the reporter revealed the spurious usage of cryptic splice acceptor and donor sites in the plasmid backbone, which were subsequently mutated (see Materials and Methods). With the corrected plasmid we performed transient transfections and confirmed the expression and circularity of the reporter generated circRNA by RT-PCR (**Figure S5B**). In addition, we validated the expression and circularity of the reporter circRNA by RNase R treatment followed by Northern blot (**Figure S5C**), and quantified RNase R resistance by RT-qPCR (**Figure S5D**).

We examined the response of our *Efnb2* backsplicing reporter (**Figure 4B**, pWT) to NOVA2 overexpression. Co-transfection of the reporter with NOVA2 led to a ~3.5-fold increase in backsplicing, whereas expression of the linear reporter RNA was unchanged (**Figure 4C**). Co-transfection of another neural-enriched splicing factor, HuD, did not alter backsplicing of circ*Efnb2* (**Figure S5E**), providing evidence of NOVA2 regulatory specificity. Together, these data demonstrate that the *Efnb2* backsplicing reporter recapitulates NOVA2 regulation of circRNAs biogenesis.

### NOVA2 regulates backsplicing of circEfnb2 via intronic YCAY motifs

To understand how NOVA2 might regulate circ*Efnb2* backsplicing, we introduced point mutations at putative NOVA2 binding sites (**Figure 4B**). Within our reporter, there were four major NOVA2 CLIP peaks, two in the downstream intron proximal to the circ*Efnb2* locus, and two preceding the 3’ splice acceptor downstream of the circRNA exon (**Figure 4A**). For RT-qPCR quantification, we normalized circRNA expression to linearly-spliced transcript expression in order to observe any relative increases in backsplicing events. Circular and linear products normalized to *Gapdh* are shown in (**Figure S5F**). We targeted the major proximal CLIP peak first, which consisted of 9 YCAY motifs. To disrupt NOVA2 binding, we mutated all 9 YCAY motifs to YAAY, since the CA dinucleotide is essential for NOVA2 recognition (69). Surprisingly, we did not observe a significant difference from pWT (**Figure 4C,** pMut1). We thus targeted the second major CLIP peak in this region consisting of two YCAY motifs, and due to potential limits of spatial resolution of the CLIP data, we considered that three adjacent YCAY motifs not identified by CLIP might also be important for backsplicing regulation. We thus mutated these additional YCAY motifs in the downstream intron. However, we still did not observe a reduction in circ*Efnb2* backsplicing when NOVA2 was overexpressed (**Figure 4C**, pMut2).

We then turned our attention to two non-clipped YCAY motifs immediately upstream of the circ*Efnb2* locus. Unexpectedly, mutations at these motifs led to a significant reduction in NOVA2-regulated backsplicing (**Figure 4C**, pMut3). Lastly, we mutated two remaining NOVA2-CLIP sites near the 3’ splice acceptor as well as two adjacent YCAY motifs not identified by CLIP. In this case, we also found significant reduction in circ*Efnb2* backsplicing compared to the WT reporter (**Figure 4C**, pMut4). This suggested that NOVA2 intronic binding on either side of a circRNA locus impacts its regulation.

### NOVA2 binding sites in circRNA flanking introns promote backsplicing

Given these reporter analysis results, we next turned to genome-wide CLIP data to investigate whether NOVA2 binding to both flanking introns is a general feature of NOVA2-regulated circRNA loci. For this analysis, we used the subset of high confidence, NOVA2-regulated circRNAs from the excitatory and inhibitory datasets (36 circRNAs for inhibitory neurons and 74 for excitatory neurons). For a non-regulated control comparison group, we used circRNAs unchanged by NOVA2 loss (*P*>0.50, FC<1) (**Supplementary File S8**). We checked for the presence of CLIP peaks in the upstream and downstream introns flanking each circRNA. We hypothesized that this robust subset would provide the best chance to identify relevant NOVA2 positional binding information. We discovered that in excitatory neurons, NOVA2 bound both flanking introns of a regulated circRNA at a significantly higher frequency than non-differentially expressed controls (*P*=0.02, Pearson’s Chi-squared test with Yates’ continuity correction) (**Figure S6A)**. In contrast, the presence of CLIP sites in just one intron (either upstream or downstream) was not significantly different between regulated circRNAs and controls. This suggests that NOVA2 intronic binding on both sides of a circRNA locus plays a role in backsplicing regulation. However, when we investigated inhibitory neurons, which had a lower sample size of regulated circRNAs, statistical significance for the same trend was not observed (**Figure S6B**).

The *Efnb2* backsplicing reporter analysis and CLIP analysis suggested that NOVA2 binding in the introns upstream and downstream of a circularizing locus promote NOVA2 regulated circRNA biogenesis. To further investigate the generality of this observation, we generated an artificial backsplicing minigene reporter, pMini, which was devoid of *Efnb2* sequences or YCAY motifs. This plasmid contains full length GFP coding sequence in the same vector backbone as our circ*Efnb2* reporter. GFP was fragmented into three artificial exons flanked by intronic sequences consisting of human *ZKscan1* reverse complementary matches (RCMs) to facilitate enhanced circRNA expression (**Figure 5A**). Existing YCAY motifs that might impact circRNA regulation were mutagenized. We confirmed that all of our pMini variants produced a single circRNA products by RT-PCR in control or NOVA2 overexpression conditions using outward facing primers (**Figure S7A**). We also confirmed the expression of the 497 nt circRNA product by Northern blot and RT-qPCR (**Figure 5B,C**), under RNase R or mock treatment conditions. As expected, RNaseR degraded the plasmid generated linear transcript (**Figure 5B,C**).

**Figure 5.**
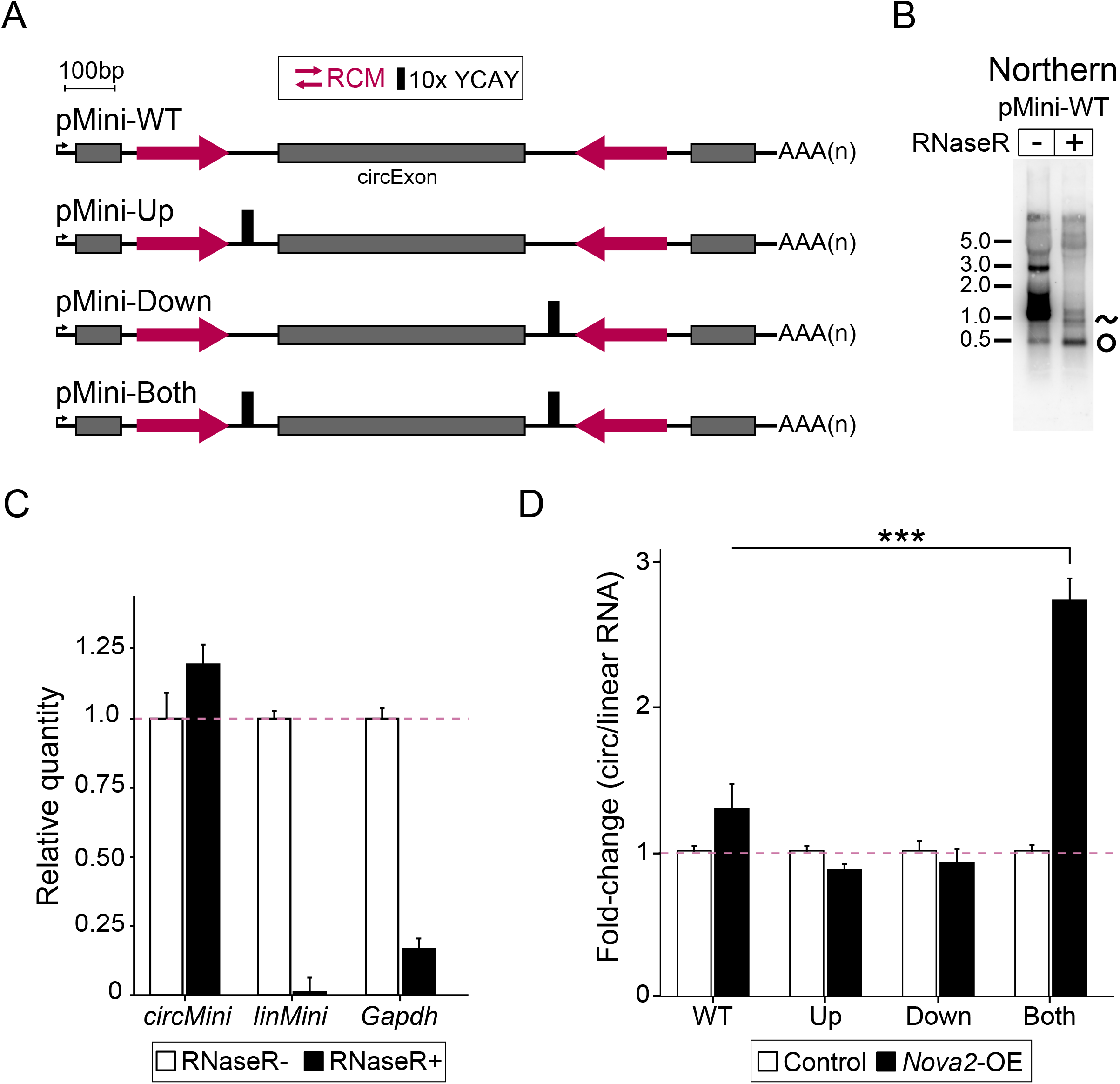
NOVA2 binding sites in both flanking introns mediate NOVA2 backsplicing. (**A**) Schematics of artificial backsplicing reporters. Exonic sequences (gray) were derived from GFP open reading frame. Reverse complementary matches (RCMs) are shown as red opposing arrows. A repeat region of 10 tandem YCAY motifs (thick black bar) was inserted into the intronic locations shown. (**B**) Northern blot using probe overlapping circularized exon of pMini reporter to detect circRNA and mRNA from transfected HEK293 cells. RNA was treated with RNase R to deplete linear transcript and enrich for circ*Mini*. (**C**) RT-qPCR expression analysis of RNase R treated RNA shown in panel B. Expression is relative to the mock RNase R condition. (**D**) RT-qPCR analysis of circ*Mini* transcript derived from pMini reporter constructs in HEK293 cells cotransfected with NOVA2 expressing plasmid (*Nova2*-OE) or empty vector control. For expression data, circ*Mini* is normalized to the linear-spliced transcript (lin*Mini*) generated by the reporter. n=3 biological replicates. Error bars are represented as mean ± SEM. \*\*\**P*<0.001, compared to pMini-WT circ*Mini* expression. Students t-test, two-tailed and unpaired.

We next introduced YCAY repeats into various intronic locations on the reporter. We introduced a 10x YCAY repeat into the WT vector (pMini-WT) ~50 bp upstream of the backspliced donor site (pMini-Up) or ~50 bp downstream of the backspliced acceptor site (pMini-Down), and in both locations (pMini-Both). For RT-qPCR quantification, we normalized circRNA expression to linearly-spliced transcript expression, as in (**Figure 4**). Circular and linear products normalized to *Gapdh* are shown in (**Figure S7B)**. Similar to pWT, introduction of YCAY repeats into the upstream intron only did not increase the ratio of circRNA to linear RNA expression (**Figure 5D**). A similar result was observed when YCAY repeats were inserted into the downstream intron only (**Figure 5D**). Finally, we tested the impact of placing NOVA2 binding sites both upstream and downstream of the circularizing exon (pMini-Both). Remarkably, we found that for this reporter, NOVA2 co-transfection led to a nearly 3-fold increase in circRNA/linear RNA ratio (**Figure 5D**, pMini-Both). Thus, similar to our circ*Efnb2* reporter, and in accordance with CLIP data from excitatory neurons, our results show that the presence of NOVA2 binding sites in both introns impacts backsplicing.

## DISCUSSION

Here we identify NOVA2 as a regulator of circRNA biogenesis in neurons. We found that within the mouse embryonic cortex, loss of NOVA2 globally reduced circRNA expression, and that this reduction was largely independent from mRNA expression changes of the host gene. This effect of global circRNA reduction upon NOVA2 loss was even more pronounced when a sorted neuron population was analyzed. We found that NOVA2-regulated circRNAs within each cell-type were largely distinct, despite overlapping expression patterns. To investigate the *cis*-elements involved in circRNA regulation by NOVA2, we focused on a conserved and abundant circRNA from the *Efnb2* gene. Using backsplicing reporter analysis we demonstrated that intronic YCAY sequences both upstream and downstream of the circRNA locus was important for NOVA2-regulation. CLIP analysis in excitatory neurons provided support for this finding.

CircRNAs are typically expressed at a low level compared to their linear counterparts. For most accepted analysis pipelines, only BSJ reads are used for quantification, making differential expression analysis problematic. Thus, we applied multiple validated pipelines to quantify circRNA expression genome-wide (55,70), and performed extensive validation of differential expression trends using RT-qPCR (**Figure 1E,F**). Investigating the sorted neuron datasets more closely, we found that NOVA2 appeared to regulate circRNAs in a cell-type specific manner (**Figure S4A**), similar to what has been previously shown for linear alternative splicing (50). Additional genome-wide analyses using library preparation methods that enhance read depth specifically for circRNAs are warranted to provide more conclusive support for this finding. On a similar front, our global analysis of how NOVA2 CLIP peaks correlated with NOVA2 regulated circRNAs could be improved by having more accurate circRNA expression quantification. There was a very low number of high-confidence NOVA2-regulated circRNAs in the Gad2 dataset (only 36)– it is possible that with greater read depth we would identify more regulated circRNAs and obtain better insight into the genome-wide features of circRNA regulation by NOVA2.

We chose circ*Efnb2*, a single-exon circRNA conserved from mouse to human, to investigate what *cis*-elements control NOVA2-regulation of circRNAs biogenesis. Using backsplicing reporter analysis we found that YCAY motifs on either side of the circularizing locus were important for generating circ*Efnb2* (**Figure 4C**). One of the important motifs identified was located in the intronic region preceding the 3’ splice-acceptor of the exon downstream of the circRNAs locus. This suggests that NOVA2 binding in this region far away from the circRNAs promotes backsplicing. A caveat of this interpretation is that several kb of the intron could not be included in our backsplicing reporter (**Figure 4A**). We discovered that two YCAY motifs upstream of circ*Efnb2* were important for regulation of backsplicing, even though they lacked NOVA2-CLIP support (**Figure 4C**). This was somewhat surprising and suggests that the CLIP datasets might have limited utility in predicting binding sites important for backsplicing. On the other hand, this result could reflect an inherent limitation of cell culture systems for recapitulating neuronal circRNAs regulation patterns. Performing mutagenesis of the intronic YCAY motifs at the endogenous *Efnb2* locus in ES-derived neurons or in mice with CRISPR genome-editing could provide more conclusive support.

To investigate NOVA2 regulation more generally, we constructed an artificial backsplicing vector, pMini, which was devoid of NOVA2 binding sites (**Figure 5A**). We introduced YCAY clusters into intronic regions upstream and downstream of the circularizing exon based on findings from the circ*Efnb2* reporter showing that binding sites on either intron were important. We found that NOVA2-induced backsplicing in th pMini reporter required the presence of YCAY clusters on both flanking introns. This result is analogous to what was previously observed for the RBP *quaking* (QKI) (38). In that study, QKI-regulated backsplicing from a reporter was found to be dependent on QKI binding sites in both upstream and downstream introns (38). QKI has been shown to self-dimerize (71). It has been hypothesized that QKI dimerization could thus be important for backsplicing regulation, although direct support for such a mechanism is currently lacking. Interestingly, NOVA proteins can also dimerize (72). Perhaps similar mechanisms of dimerization are involved in backsplicing regulation by NOVA2.

*Nova2*-KO mice display a host of degenerative brain phenotypes which have been attributed to deregulation of linear alternative splicing (13,49,50). Hundreds of circRNAs were found here to be differentially regulated by *Nova2*. Could reduced levels of circRNAs such as circ*Efnb2* contribute to the neurodevelopmental defects of *Nova2*-KO mice? There are many possible ways NOVA2-regulated circRNAs could impact neural development, given the different ways circRNAs impact gene regulation. For example, some circRNAs to travel to synapses and act as scaffolds for various RBPs, whereas others regulate the transcriptional activity of genes in the nucleus (30,74,75). Some circRNAs have been recently associated with neurological defects in mice (34) and humans (36,76). Despite technical challenges, several recent studies have demonstrated the feasibility of both targeting circRNAs using RNAi (35,36,77) and deleting intronic RCMs using CRISPR to reduce or eliminate circRNA expression (75). Moving forward, it will be interesting to assess the role of NOVA2-regulated circRNAs such as circ*Efnb2* in neural development using such approaches.

### SOFTWARE AVAILABILITY

CIRI2 is an open source, freely available tool for access at Source Forge (https://sourceforge.net/projects/ciri/files/CIRI2/)

HISAT2 is an open source, freely available tool for access at Johns Hopkins University (https://ccb.jhu.edu/software/hisat2/manual.shtml)

STAR is an open source, freely available tool in the GitHub repository (https://github.com/alexdobin/STAR)

DCC/CircTest is an open source, freely available tool in the GitHub repository (https://github.com/dieterich-lab/DCC) and (https://github.com/dieterich-lab/CircTest)

rMATS is an open source, freely available tool at Source Forge (http://rnaseq-mats.sourceforge.net)

IGV is freely available tool at the Broad Institute (https://software.broadinstitute.org/software/igv/)

## SUPPLEMENTARY DATA

**Supplementary File S1:** RNA-seq datasets used in this study

**Supplementary File S2:** CIRI2 circRNA expression *Nova1*-KO vs WT E18.5 cortex

**Supplementary File S3:** CIRI2 circRNA expression *Nova2*-KO vs WT E18.5 cortex

**Supplementary File S4:** CIRI2 circRNA expression *Nova2*-KO vs WT E18.5 Emx1+ neurons

**Supplementary File S5:** CIRI2 circRNA expression *Nova2*-KO vs WT E18.5 Gad2+ neurons

**Supplementary File S6:** DCC/CircTest host-mRNA and circRNA expression *Nova2*^−/−^ vs WT E18.5 Emx1/Gad2+ neurons

**Supplementary File S7:** rMATS AS events and DE-circRNA overlap *Nova2*-KO vs WT E18.5 Emx1+/Gad2+ neurons

**Supplementary File S8:** Overlap between CIRI2/CircTest results *Nova2*^−/−^ vs WT E18.5 Emx1/Gad2+ neurons

**Supplementary File S9:** Oligonucleotides primers/probes used for RT-qPCR, RT-PCR, and Northern blot

## ACKNOWLEDGEMENT

We thank Dr. Zhe Chen (University of Minnesota) for sharing plasmids. We thank Nora Perrone-Bizzozero (University of New Mexico) for sharing the HuD plasmid. We thank A. van der Linden, Z. Chen, Z. Zhang, and B. Bae for helpful feedback and discussion on the manuscript.

