## Supplementary material for "NOVA2 regulates neural circRNA biogenesis": Figure S

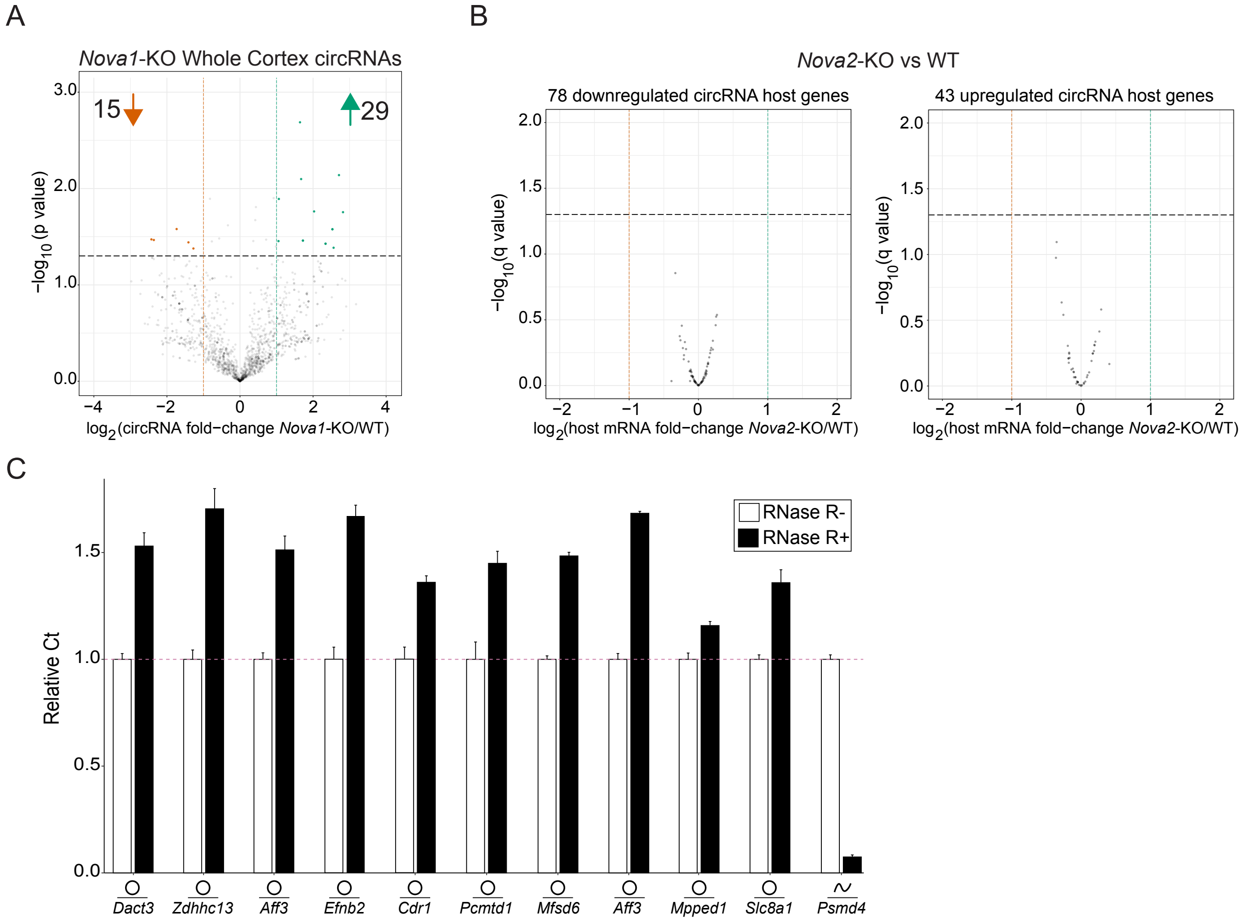


**Supplementary Figure 1. NOVA1 circRNA regulation, NOVA2-regulated circRNA host gene mRNA expression, and RNase R resistance of circRNAs.** (**A**) Volcano plot of circRNA expression in *Nova1*-KO vs. WT mouse cortex ($log_2FC>1$, $P<0.05$). (**B**) Host-gene linear RNA expression of downregulated circRNAs (*left panel*) and upregulated circRNAs (*right panel*) identified by CIRI2 are not differentially expressed in *Nova2*-KO cortex. *P* values were obtained by the Wald test and corrected for multiple hypothesis testing using the Benjamini and Hochberg method (DESeq2 default settings). (**C**) RT-qPCR relative expression analysis of 10 circRNAs (shown in **Figure 1E-F**) and one linear RNA control (*Psmd4*). Total RNA was treated with RNase R or mock control prior to cDNA synthesis. Expression is relative to the mock RNase R condition (RNase R-). Error bars are represented as the mean ± SEM.

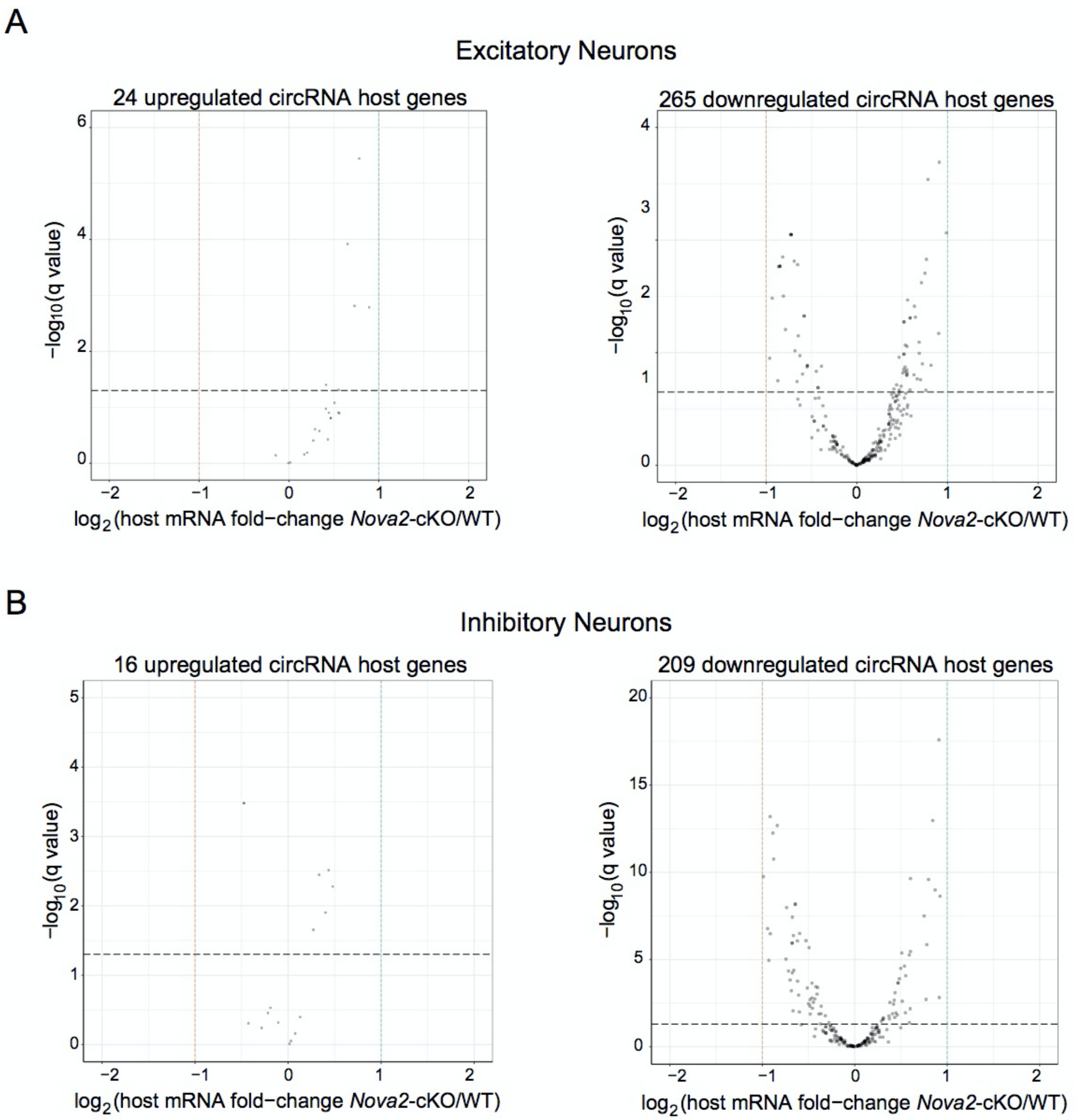


**Supplementary Figure 2. NOVA2-regulated circRNA host gene mRNAs are not differentially expressed in excitatory or inhibitory neuron datasets.** (**A**) Host gene mRNA expression of upregulated circRNAs (*left panel*) and downregulated circRNAs (*right panel)* from *Nova2*-KO excitatory neurons are not significantly differentially expressed compared to WT samples ($Log_2FC>1$, $q<0.05$). (**B**) The same analysis performed for inhibitory neurons. *P* values for each dataset were obtained using the same approach as in Figure S1B.

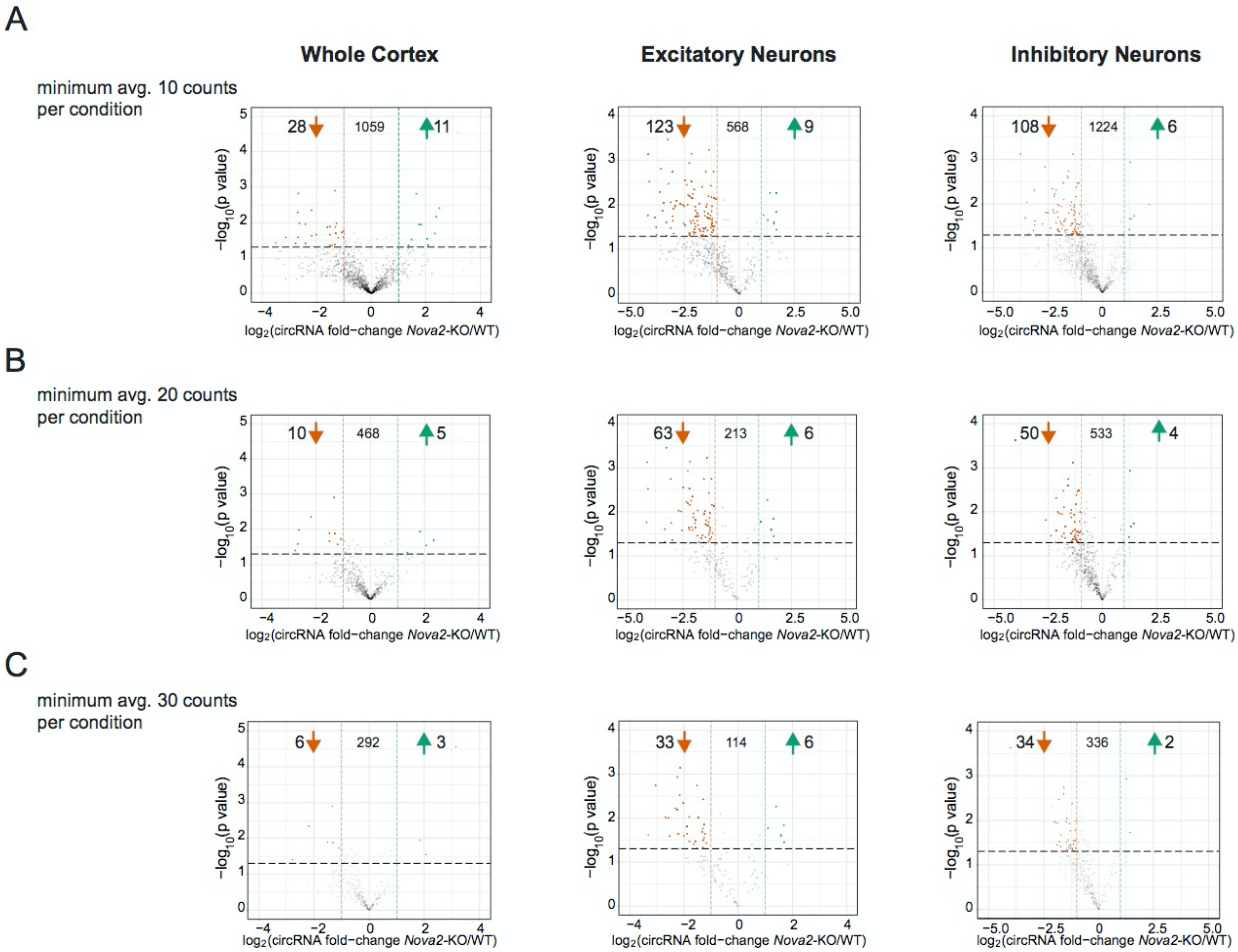


**Supplementary Figure 3. Reduced circRNA expression by NOVA2 is consistent among increasing minimum BSJ count thresholds.** Volcano plots of circRNAs identified by CIRI2 are regulated among WT vs *Nova2*-KO whole cortex, excitatory neuron, and inhibitory neuron RNA-seq datasets with minimum averages of (**A**) 10, (**B**) 20, and (**C**) 30 BSJ read count thresholds applied. The number of circRNAs passing each read count threshold are shown in the center of each plot. Significance reflects a $log_2FC>1$ and $P<0.05$.

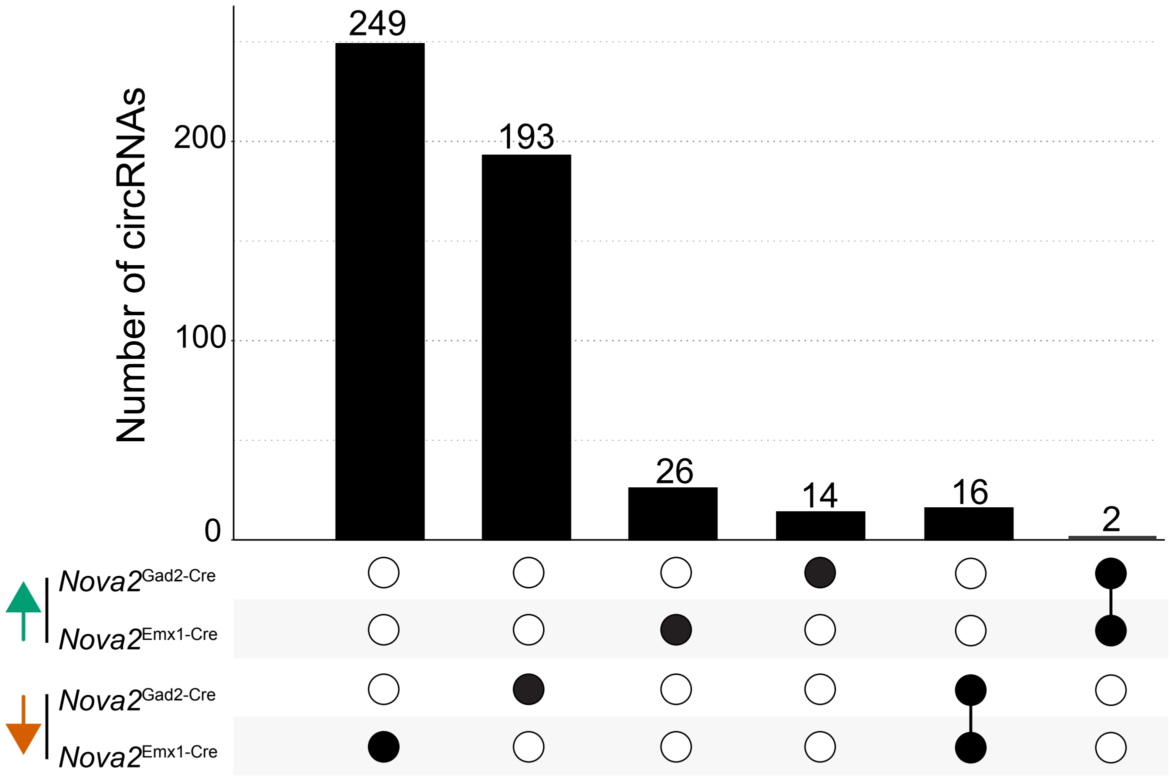


**Supplementary Figure 4. NOVA2-regulated circRNAs in neurons exhibit cell-type specificity.** (**A**) Number of differentially expressed circRNAs unique or overlapping in excitatory and inhibitory neuron datasets. Green up arrow represents *Nova2*-cKO upregulated circRNAs. Orange down arrow represents *Nova2*-cKO downregulated circRNAs. CircRNAs were identified using CIRI2.

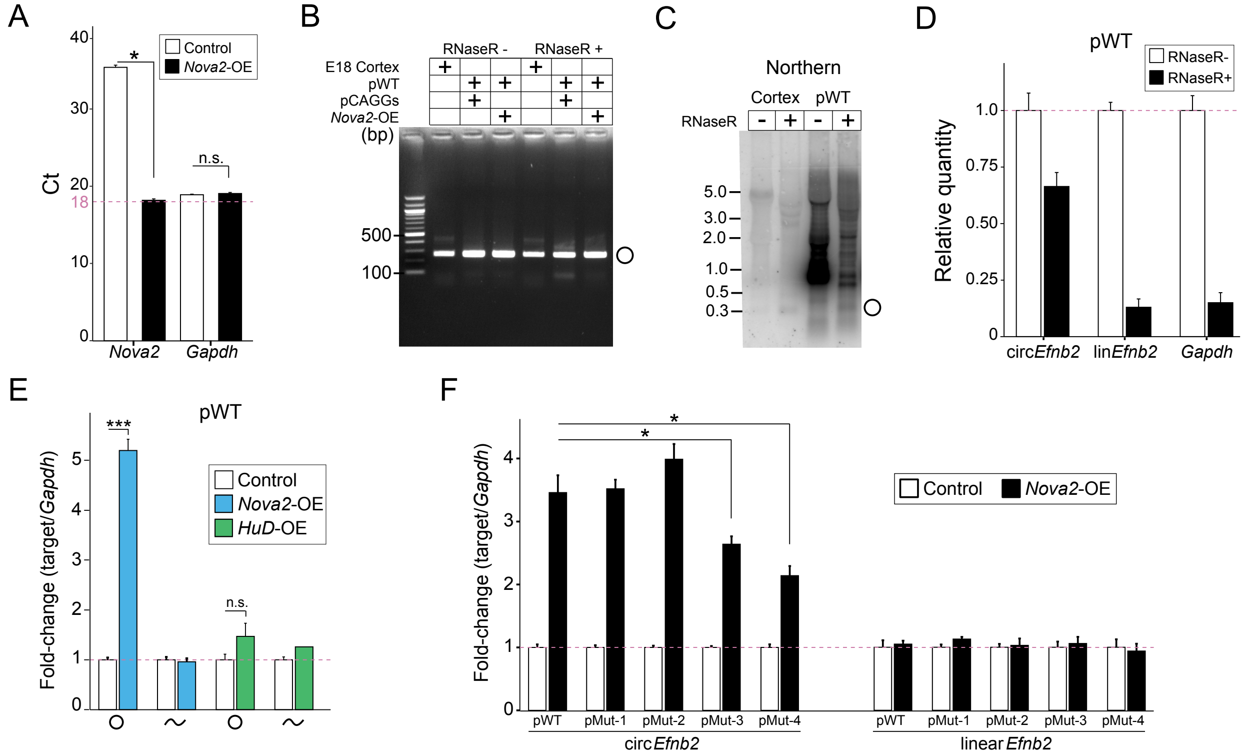


**Supplementary Figure 5. Circ*Efnb2* backsplicing reporter generates a single, RNase R resistant, circ*Efnb2* transcript responsive to NOVA2**. (**A**) RT-qPCR relative expression of *Nova2* and *Gapdh* transcripts in HEK293 cells pre- and post-transfection with NOVA2 expression (*Nova2*-OE) plasmid (n=3). (**B**) RT-PCR analysis of circ*Efnb2* transcript from pWT backsplicing reporter with or without RNase R treatment and pre- and post-transfection with NOVA2 expression plasmid in HEK293 cells. Outward facing primers were used to capture a 267 bp product. (**C**) Northern blot analysis utilizing a dsDNA probe targeting the circular and linear transcripts of *Efnb2* gene extracted from mouse cortex or HEK293 cells transfected with the pWT reporter. Circ*Efnb2* signal was enriched post-RNase R treatment while linear transcripts were diminished. (**D**) RT-qPCR relative expression analysis of circ*Efnb2* and linear *Efnb2* transcripts generated from pWT vector. RNA used to generate cDNA for plot is the same RNA used for Northern blot in panel B. The circ*Efnb2* transcript is more resistant to RNase R treatment relative to the linear *Efnb2* transcript and endogenous *Gapdh* control gene. (**E**) RT-qPCR expression analysis of circ*Efnb2* and lin*Efnb2* generated from pWT reporter pre- and post-transfection with either NOVA2 or HuD expression (*HuD*-OE) vectors. Expression is normalized to *Gapdh* (n=3). (**F**) RT-qPCR expression analysis for circ*Efnb2* (*left*) or linear *Efnb2* (*right*) transcripts generated from pWT and mutated reporter variants pre- and post-transfection with NOVA2 expression vector. Data is related to **Figure 4C**. Expression is normalized to *Gapdh* (n=3). For expression analyses in panel A, E, and F, Student's t-test was used for statistical significance (two-tailed, unpaired) *$P$<0.05, ***$P$<0.001, n.s., not significant.

A

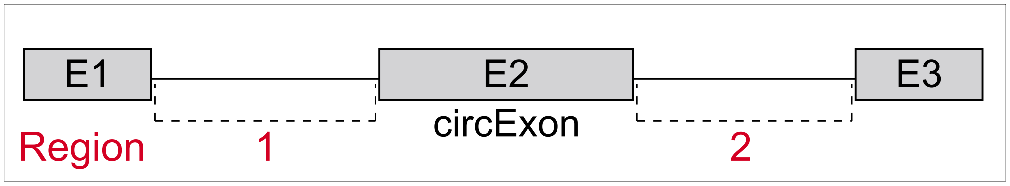


Excitatory neurons

| *Nova2*-CLIP Region | 1 | 2 | 1 or 2 | 1 and 2 |
|---|---|---|---|---|
| # of Circs (Control*) | 270 | 253 | 378 | 145 |
| # of Circs (Regulated**) | 42 | 43 | 55 | 30 |
| Percent (Control) | 50% | 47% | 70% | 26% |
| Percent (Regulated) | 57% | 58% | 74% | 41% |
| *P* value | 0.33 | 0.09 | 0.53 | 0.02 |

*540 circRNAs **74 circRNAs

B

Inhibitory neurons

| *Nova2*-CLIP Region | 1 | 2 | 1 or 2 | 1 and 2 |
|---|---|---|---|---|
| # of Circs (Control*) | 640 | 591 | 881 | 350 |
| # of Circs (Regulated**) | 18 | 18 | 23 | 13 |
| Percent (Control) | 46% | 42% | 63% | 25% |
| Percent (Regulated) | 50% | 50% | 64% | 36% |
| *P* value | 0.75 | 0.46 | 1.00 | 0.19 |

*1393 circRNAs **36 circRNAs

**Supplementary Figure 6. NOVA2-CLIP peak analysis of NOVA2-regulated circRNAs overlapping between CIRI2 and DCC/CircTest analyses.** (**A**) (*Top*) Schematic of two intronic regions flanking NOVA2-regulated circRNA loci examined for significant NOVA2-CLIP peak presence. The number of NOVA2-regulated (74) or control (540, FC<1, $P$>0.50) circRNAs overlapping an intronic NOVA2-CLIP peak within each defined region from the excitatory neuron dataset is shown below. (**B**) The same analysis was performed for 36 NOVA2-regulated circRNAs and 1393 control circRNAs from the inhibitory neuron dataset. *P* values were generated by Pearson's Chi-squared test with Yates' continuity correction.

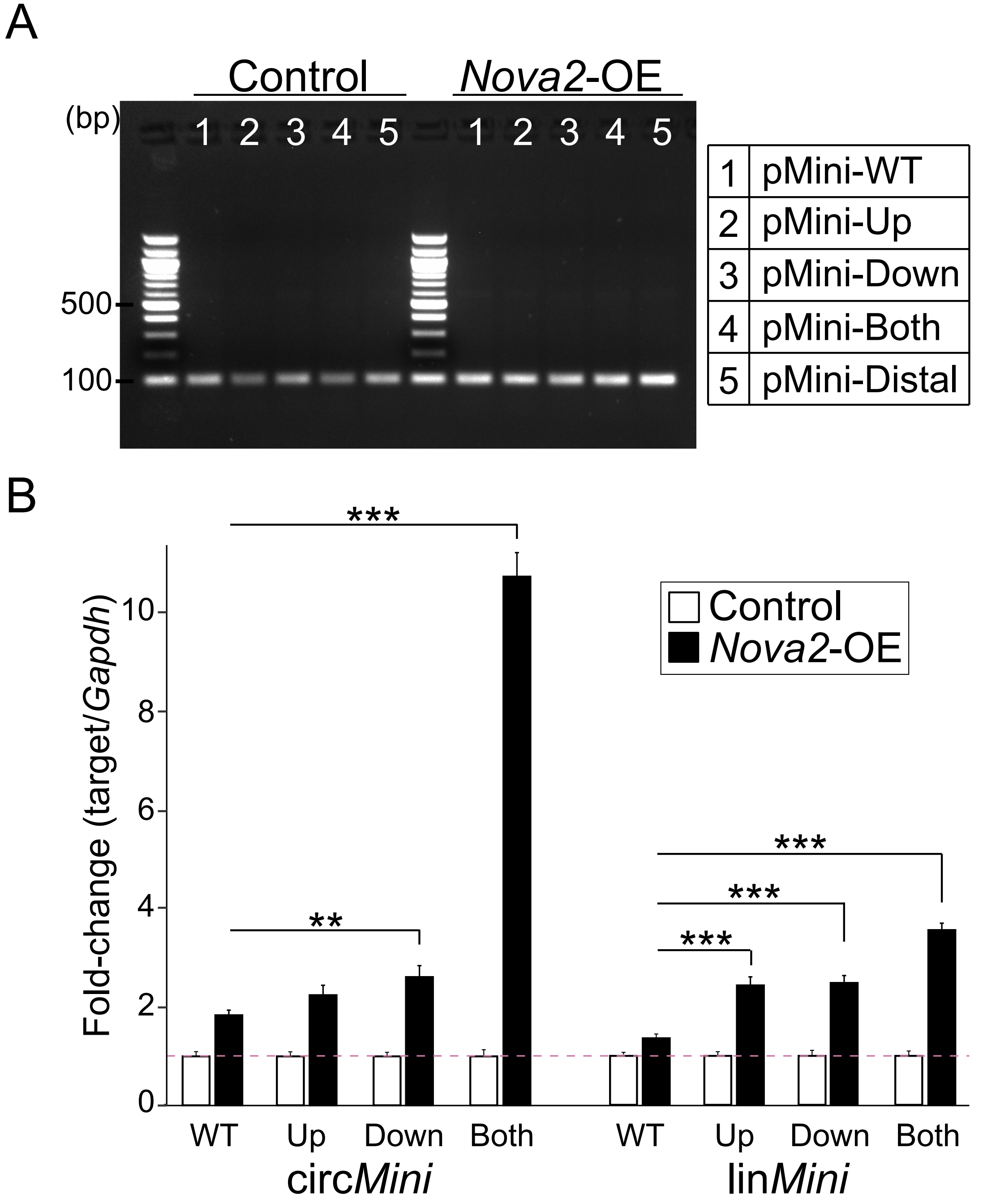


**Supplementary Figure 7. pMini backsplicing reporter circular transcript validation and quantitative expression analysis. (A**) RT-PCR analysis of circ*Mini* transcript generated from the five pMini backsplicing reporters pre- and post-transfection with NOVA2 expression (*Nova2*-OE) plasmid in HEK293 cells. Outward facing primers were used to capture a 101 bp product. (**B**) RT-qPCR expression analysis of circular (circ*Mini*) and linear (lin*Mini*) transcripts generated from the pMini backsplicing reporters in the presence or absence of NOVA2 related to **Figure 5D**. Expression is normalized to *Gapdh* (n=3). Student's t-test was used for statistical significance (two-tailed, unpaired) ***P*<0.01, ****P*<0.001.
